# Retinotopic remapping of the visual system in deaf adults

**DOI:** 10.1101/2020.01.31.923342

**Authors:** Alexandra T. Levine, Kate Yuen, André Gouws, Alex R. Wade, Antony B. Morland, Charlotte Codina, David Buckley, Heidi A. Baseler

**Author notes:** Correspondence should be addressed to: Dr. Heidi A. Baseler.

## Abstract

Deaf individuals rely on visual rather than auditory cues to detect events in the periphery, putting a greater demand on neural resources for vision. Comparing visual maps in the brains of early deaf and hearing adults, we found a redistribution of neural resources in the lateral geniculate nucleus and primary visual cortex, with larger representations of the periphery, at a cost of smaller representations of the central visual field.

## Introduction

Sound can serve as a vital cue, helping hearing people orient their gaze and attention towards events outside their central line of sight, especially in the far periphery where vision is poor. Without sound cues, deaf people must rely on vision as an ‘early warning system’ for peripheral events; indeed, numerous behavioural studies demonstrate superior visual sensitivity in early deaf individuals (for reviews see Bavelier et al., 2006; Pavani & Bottari 2012), particularly to far peripheral stimuli (Lomber et al., 2010; Buckley et al., 2010; Codina et al., 2017). Here, we provide evidence that a lifelong reliance on vision in deaf individuals can lead to a redistribution of inputs in early visual brain regions, increasing neural resources for processing stimuli in the far peripheral visual field.

Previous studies revealed differences in visually evoked cortical potentials between deaf and hearing groups, suggesting potential neural substrates for enhanced visual sensitivity (Neville & Lawson, 1987). Later studies using brain imaging and animal models indicated that deaf adults recruit auditory regions of the brain in response to visual stimuli (cross-modal plasticity) (Finney et al., 2001; Fine et al., 2005; Lomber et al., 2010; Bottari et al., 2014; Cardin et al., 2016; Shiell et al., 2016; Szwed et al., 2017). Other studies revealed neural differences in deaf individuals in higher-order visual areas, such as V5/hMT+ (Bavelier, 2001; Scott, 2014), but not in early visual areas (Fine et al., 2005). Expanded representations of intact inputs in primary sensory cortex have been demonstrated in other populations with sensory loss, e.g., amputees (Merzenich et al., 1984; Ramachandran & Rogers-Ramachandran, 2000) and those with early partial sight loss (Baseler et al., 2002; Legge 2011). Early visual reorganisation is also evident in other populations with lifelong differences in sensory behaviour, e.g. amblyopes (Conner et al., 2007) and children with autistic spectrum disorder (ASD) (Frey et al., 2013). Moreover, structural changes as early in the visual pathways as the retina have been correlated with enhanced far peripheral sensitivity in deaf adults, leading to the possibility that early visual structures downstream may also be affected (Codina et al., 2011). Previous studies evaluating early visual cortex have found either no difference between deaf and hearing cortex (Fine et al., 2005) or thinner visual cortex representing the peripheral visual field (Smittenaar et al., 2016). However, these studies were confined to comparisons within a limited visual field (< 37.5°), rather than the far periphery where differences in visual sensitivity are most prominent and potentially beneficial (Buckley et al., 2010; Codina et al., 2017).

## Results

We mapped visual field representations including the far periphery in primary visual cortex and the lateral geniculate nucleus in a group of early, profoundly deaf adults (N=16) and a group of hearing age-matched controls (N=16) using functional magnetic resonance imaging. Wide-field retinotopic mapping stimuli were used to identify primary visual cortex (V1) and map visual field eccentricity out to 72° (Fig. 1a). In order to compare eccentricity representations, V1 was divided into three roughly equal-sized sub-regions of interest – central (0-15°); mid-peripheral (15-39°) and far-peripheral (39-72°) (Fig. 1a; see methods for more details). Left and right lateral geniculate nuclei (post-retinal visual relay structure in the thalamus) were identified anatomically in each participant using structural (proton density) images (Fujita et al., 2001; Fig. 1b).

**Figure 1.**
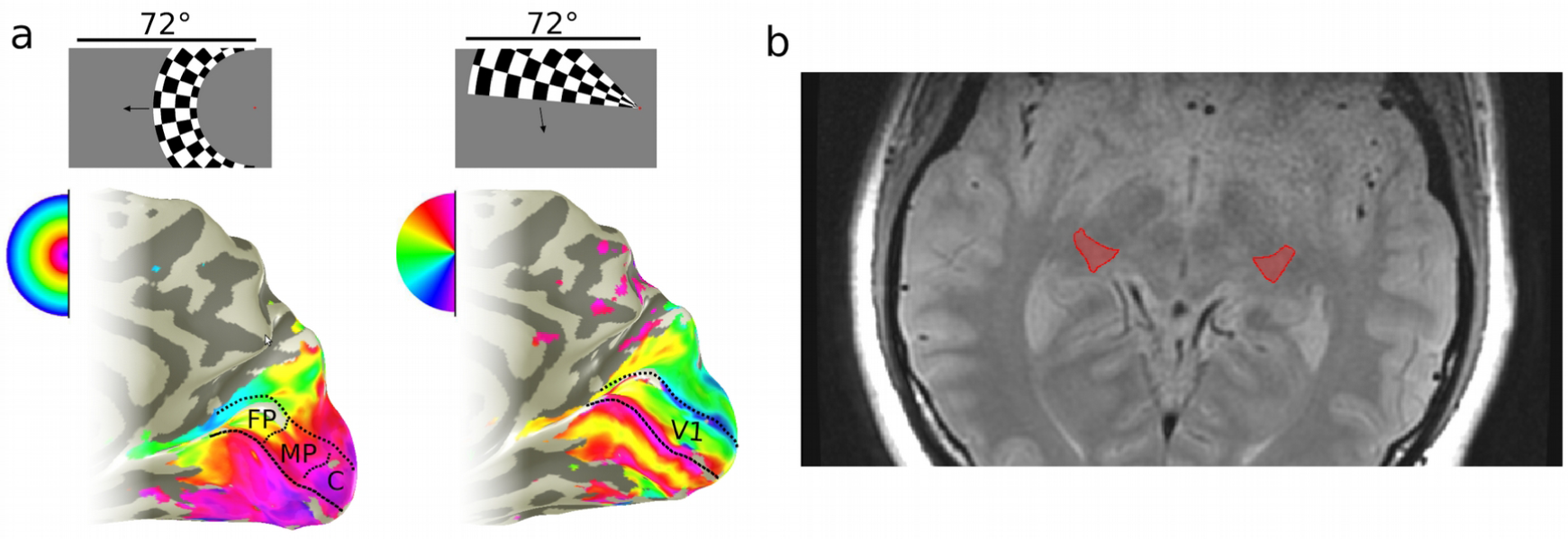
Regions of interest in primary visual cortex (V1) and the lateral geniculate nucleus (LGN) (a) Visual mapping in primary visual cortex (V1). Visual fMRI response maps to expanding ring stimulus (left), and rotating wedge stimulus (right) superimposed on the medial surface of an individual inflated right occipital lobe. Example stimuli shown above. Semicircular key indicates the stimulus position in the visual field maximally activating each part of visual cortex, in false colour. Three regions of interest in V1 marked schematically on left image representing the central (C, 0-15°); mid-peripheral (MP, 15-39°), and far-peripheral (FP, 39-72°) visual field. (b) Single axial slice of a proton density scan in one participant. Left and right lateral geniculate nuclei (LGN) are indicated in this slice in red.

First, we compared the total cortical **volume** of primary visual cortex. We found no difference between deaf and hearing groups (Fig. 2a) (t(28.19)= -0.154, p = .878), in agreement with a previous study that compared visual map size between deaf and hearing adults within a smaller, central visual field representation (+/-20°; Fine et al., 2005).

**Figure 2.**
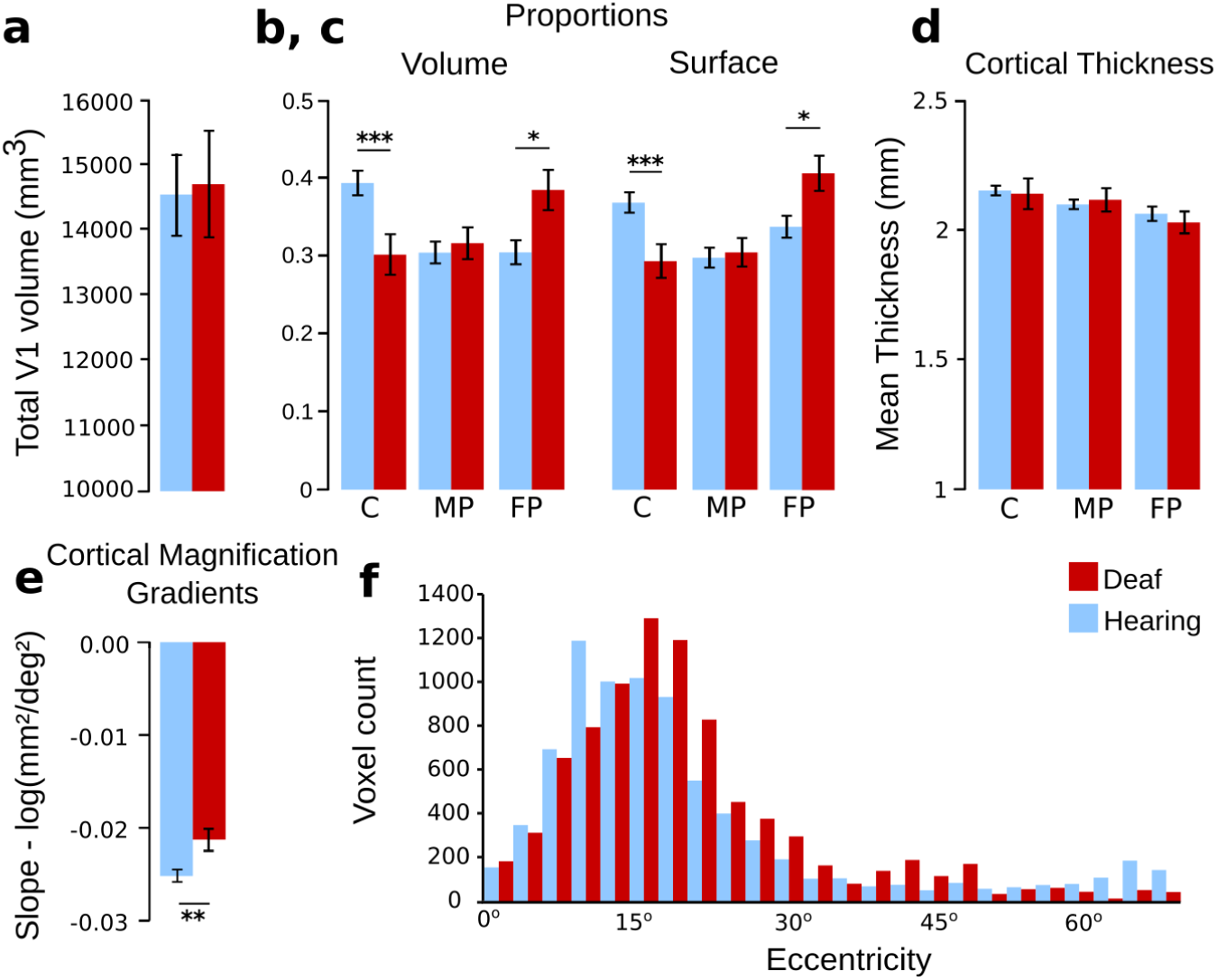
V1 total cortical volume, mean cortical thickness, proportion of volume and surface area in V1, cortical magnification gradients and LGN voxel responsiveness to the expanding ring stimulus. (a) Mean total V1 cortical volume based on wide field retinotopic mapping. Mean cortical volume (b) and cortical surface (c) proportions in V1 sub-regions. (d) Mean cortical thickness of V1 areas, grouped by representation of visual field eccentricity. (e) Mean cortical magnification gradients of V1. (e) Preferred eccentricity for voxels in the lateral geniculate nucleus (LGN). Sub-regions represent the central (C, 0-15°); mid-peripheral (MP, 15-39°), and far-peripheral (FP, 39-72°) visual field. Error bars indicate +/- one standard error of the mean. ***p<0.001, **p<0.01, *p<0.05.

To correct for individual differences in overall V1 size, we compared the **proportional volume** of the three visual field representations in V1 between deaf and hearing groups. A mixed factorial analysis of variance (ANOVA) revealed a significant interaction between sub-region and group (F(1,30)= 7.608, p = .002, η2= .923), indicating that the volume of V1 is distributed differently across the three visual field representations between deaf and hearing groups (Fig. 2b). Between-group comparisons indicated a significantly larger central representation in hearing participants (t(23.22) = 2.98, p = .007), no significant difference in the mid-peripheral representation (t(26.83) = -0.452, p = .655), and a significantly larger far-peripheral representation in deaf participants (t(24.93) = -2.62, p = .015: Fig. 2b). This result suggests that primary sensory cortex accommodates neuroplasticity by trading off central and far peripheral representations, rather than increasing in overall size.

As differences in cortical volume can be affected by either changes in cortical thickness or surface area or both, we compared both measures separately between groups. We found no difference in overall **mean cortical thickness** (all of V1) between deaf and hearing groups (ANOVA; Group, F(1,30) = .037, p = .849, η2 = .054; see Fig. 2d). There was a significant difference in thickness between eccentricity representations across both groups (Region, F(1.589,47.674) = 14.145, p < .001, η2 = .992), supporting the well-documented decrease in thickness in all human brains between the occipital pole (representing the central visual field) and the calcarine sulcus (representing the periphery) (Griffis et al., 2016). Importantly, however, there was no significant interaction between region and group, suggesting cortical thickness does not account for the volume differences found above (Region*Group: F(1.589,47.674)= .923, p = .384, η2= .183). This is in contrast to previous studies showing either decreased (Smittenaar et al. 2016) or increased cortical thickness in the calcarine sulcus of deaf individuals (Allen et al., 2013; Penicaud et al., 2013).

**Cortical surface area** of each eccentricity representation was compared in terms of its proportion of V1 to correct for individual differences in overall V1 surface area (Fig. 2c). The ANOVA showed a significant interaction between the proportion of cortical surface of the three V1 ROIs and group (F(1,30)= 7.608, p = .002, η2= .923), as seen in the volume measure. Between-group comparisons indicated a significantly larger central representation in hearing participants (t(21.53) = 2.93, p = .008, d = .10), no significant difference in the mid-peripheral representation (t(27.42) = -0.244, p = .809, d = .08), and a significantly larger far-peripheral representation in deaf participants (t(25.97) = -2.50, p = .019, d= .91: Fig. 2c). This suggests that volumetric differences between groups were related to surface area changes rather than cortical thickness.

The distribution of neural resources allocated to processing different visual field locations is conventionally expressed as the areal **cortical magnification** function (cortical area per unit visual field area), which decreases exponentially with eccentricity in V1 (Dougherty et al., 2003; Adams & Horton, 2003). To compare the cortical magnification function in V1 between deaf and hearing groups, we calculated the cortical area per unit visual field area represented by each eccentricity region in each participant. Logarithmic values were plotted as a function of eccentricity and fit with a linear regression line to estimate gradients (see methods for more details). A comparison of gradients revealed they were significantly shallower in deaf than hearing participants (t(23.56) = -2.96, p = .007, d = 2.67), illustrating a bias towards larger far peripheral representations at the expense of smaller central representations in primary visual cortex in deaf relative to hearing individuals (Fig. 2e).

Next, we asked whether the redistribution of visual field resources in deaf participants was unique to visual cortex. Previous evidence that projections from peripheral retina are thicker in deaf than hearing adults suggests that cortical differences might be inherited from an earlier stage in the visual pathways (Codina et al., 2011). If so, a peripheral bias in visual field representations might also be detectable in the **lateral geniculate nucleus** (LGN), the thalamic relay structure that receives inputs from the retina and is the main source of projections to V1. To examine this, LGN voxels from each participant were categorised according to preferred eccentricity (stimulus eccentricity producing maximal functional response). Cumulative histograms were plotted as a function of eccentricity for each group (Fig. 2f). Distributions were significantly different between groups (two-sample Kolmogorov-Smirnov test, D = 0.094, p < 0.001). The deaf group showed a higher median (18.25) compared to the hearing (16.23). Both groups are positively skewed, with the hearing showing a more positive skew (1.62), indicating a higher preference towards lower eccentricities, when compared to the deaf (skew = 1.44). Overall, the LGN data indicates a response bias towards the peripheral visual field in the deaf group relative to the hearing group. This suggests that the redistribution of visual field representations found in primary visual cortex may be inherited in part from earlier structures, such as the LGN.

## Discussion

Our results provide the first evidence of compensatory plasticity in the LGN and V1 as a consequence of lifelong auditory deprivation. We reveal a redistribution of neural resources in early deaf individuals, with a larger cortical surface representation of the periphery (>39°), at a cost of smaller representations of the central visual field (<15°).

This central/peripheral trade-off in neural resources may explain behavioural differences reported between deaf and hearing individuals, particularly increased visual sensitivity in the far peripheral visual field (Buckley et al., 2010; Pavani & Bottari, 2010; Codina et al., 2017; Almeida et al., 2015). Although many studies show no perceptual differences between deaf and hearing adults within the central visual field (Bosworth and Dobkins 1999; Brozinsky and Bavelier 2004; Stevens and Neville 2006), some disadvantages have been noted in deaf adults (Proksch & Bavelier, 2002; Parasnis, Samar & Berent, 2003; Samar & Berger, 2017), possibly explained by the reduction of neural resources we find in the central visual field representations. Furthermore, there is evidence of a central/peripheral trade-off in visual attention of deaf individuals (Proksch & Bavelier, 2002; Parasnis, Samar & Berent, 2003) and responses measured in visual motion area V5/hMT+ (Bavelier et al., 2001). Few studies find a relationship between structure and visual performance in hearing (Duncan & Boynton, 2003) and deaf participants (Codina et al., 2011; Smittenaar et al., 2016; Shiell et al., 2016). However, it is likely that a combination of factors contribute to improved visual performance in deaf individuals, including the well-established crossmodal recruitment of auditory brain regions (Lomber et al., 2010; Shiell et al., 2016). Nevertheless, it appears that neural plasticity in the deaf is not limited to cortical regions deprived of input, i.e. auditory cortex, but is also present unimodally within the visual system as well.

Most of our deaf participants were fluent and frequent users of British Sign Language for communication, leading to the possibility that visual experience through signing might contribute to the neural changes observed. Although we observed the same pattern of central/peripheral differences between deaf and hearing groups in both hemispheres, they were most pronounced in the left hemisphere. This might reflect the right visual field advantage reported in deaf (and hearing) signers (Bosworth & Dobkins, 1999, 2002; Neville & Lawson, 1987b; Codina et al., 2017) compared to hearing non-signers. However, sign language perception is primarily confined within the central/mid-peripheral visual field rather than the far periphery (Stoll et al., 2018). Moreover, previous studies comparing deaf and hearing native sign language users suggest that plastic changes are primarily due to sensory deprivation, with only modest or no changes with sign language use (Bavelier et al., 2006; Cardin et al., 2015; Codina et al., 2017; Stoll et al., 2018).

Differences in neural resources within the LGN and V1 may partly reflect upstream differences in projections from the retina. Codina et al. (2011) reported that the retinal nerve fibre layer containing ganglion cell axons from peripheral retina is thicker in deaf participants, whilst the region containing central projections is thicker in hearing participants. As the size of the optic tract, LGN and visual cortex are correlated within individuals (Andrews et al., 1997), these differences can lead to changes in downstream projection areas. Our novel findings in the LGN suggest that increased cortical representation is at least in part mediated by feedforward connections. Parallel studies in the somatosensory system also indicate that thalamic projections can influence cortical plasticity further along the processing pathway (Jain et al., 2008). However, differences in the visual field distribution of electrophysiological responses from the retina and cortex in deaf and hearing groups suggest that additional modifications may occur at the level of the cortex (Baseler et al., 2013). Finally, changes in the peripheral representation in V1 might also reflect input from direct crossmodal connections from auditory and superior temporal regions to calcarine cortex (Falchier et al., 2002; Rockland & Ojima 2003; Beer et al., 2011; Shiell et al., 2014).

For the first time, this study demonstrates that lifelong reliance on peripheral vision in congenitally, profoundly deaf individuals leads to a redistribution in neuronal resources, favouring the far periphery, at the expense of representations of the central field, and that evidence of this reorganization can be seen at multiple levels of the ascending visual hierarchy. This compensatory plasticity in deaf adults was present at the level of the lateral geniculate nucleus of the thalamus, as well as early visual cortex. These results contribute to the general debate around the stability and plasticity of early visual maps. Changes in visual maps have been demonstrated when central retinal input is absent from birth (Baseler et al., 2002). However, when retinal input is lost later in life, the extent and nature of reorganisation is less clear (Baseler et al., 2011; Dilks et al., 2008; Masuda et al., 2008).

Taken together, our findings support the notion that greater reliance on the far peripheral visual field in congenitally, profoundly deaf individuals drives neural plasticity within the early visual system.

## Acknowledgements

The authors would like to thank Shradha Billawa, Laura Bridge, Sally Clausen, Eleanor Cole, Vera Wang, Shanelle Canavan, Lucy Spencer, Charlotte Campbell for their contributions in data collection, and the staff at The York Neuroimaging Centre for all their support during the course of this project. Furthermore, we would like to thank Dr Mark Hymers and Dr Edward Silson for useful discussion of the study.

## Author contributions

Alexandra T. Levine: conceptualization, methodology, software, formal analysis, writing of original draft, visualization

Kate Yuen: formal analysis, writing - review and editing

André D. Gouws: methodology, formal analysis

Alex R. Wade: methodology, formal analysis, supervision

Antony B. Morland: conceptualization, resources, supervision

Charlotte Codina, David Buckley: writing - review and editing, resources

Heidi A. Baseler: conceptualization, methodology, writing - review and editing, resources, supervision, funding acquisition, project administration, visualization

## Methods

### Participants

The study included 32 participants: 16 early deaf individuals (mean age=34.13, range=20-48 years, 5 females) and 16 hearing individuals (mean age=29.61, range=20-48 years, 5 females). There was no significant difference in age between the groups (t(30)=1.291, p=.207). All participants had normal or corrected-to-normal vision and gave informed consent in accordance with the Declaration of Helsinki. A British Sign Language (BSL) interpreter was present throughout the sessions for BSL-speaking deaf participants to ensure they understood the instructions and purpose of the study and to answer any questions. Each deaf participant also filled out a brief questionnaire regarding the known aetiology of deafness (Table 1). All deaf participants reported severe to profound hearing loss in both ears (>70db) since infancy (<3 years). The study was approved by The York Neuroimaging Centre Research Governance Committee.

**Table 1.**
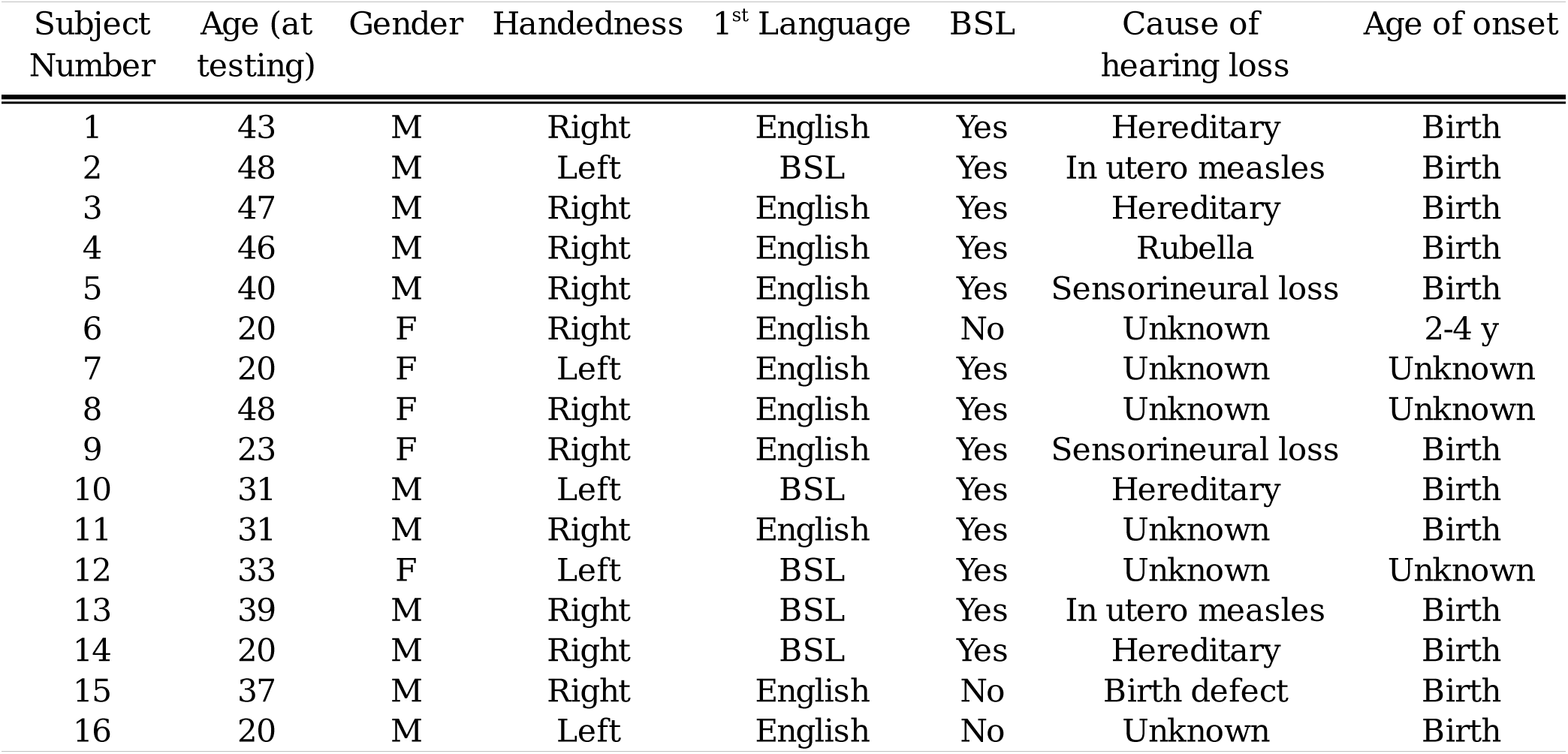
A list of all deaf participants included in the current study.

| Subject Number | Age (at testing) | Gender | Handedness | 1<sup>st</sup> Language | BSL | Cause of hearing loss | Age of onset |
| --- | --- | --- | --- | --- | --- | --- | --- |
| 1 | 43 | M | Right | English | Yes | Hereditary | Birth |
| 2 | 48 | M | Left | BSL | Yes | In utero measles | Birth |
| 3 | 47 | M | Right | English | Yes | Hereditary | Birth |
| 4 | 46 | M | Right | English | Yes | Rubella | Birth |
| 5 | 40 | M | Right | English | Yes | Sensorineural loss | Birth |
| 6 | 20 | F | Right | English | No | Unknown | 2-4 y |
| 7 | 20 | F | Left | English | Yes | Unknown | Unknown |
| 8 | 48 | F | Right | English | Yes | Unknown | Unknown |
| 9 | 23 | F | Right | English | Yes | Sensorineural loss | Birth |
| 10 | 31 | M | Left | BSL | Yes | Hereditary | Birth |
| 11 | 31 | M | Right | English | Yes | Unknown | Birth |
| 12 | 33 | F | Left | BSL | Yes | Unknown | Unknown |
| 13 | 39 | M | Right | BSL | Yes | In utero measles | Birth |
| 14 | 20 | M | Right | BSL | Yes | Hereditary | Birth |
| 15 | 37 | M | Right | English | No | Birth defect | Birth |
| 16 | 20 | M | Left | English | No | Unknown | Birth |

### MRI Protocol

MRI data were acquired using a 16 channel posterior brain array coil (Nova Medical) in a GE 3 Tesla Signa Excite HD scanner at the York Neuroimaging Centre. High-resolution structural data of the entire brain were acquired with high resolution T1-weighted isotropic scans, (TR, 8 ms; TE, 3 ms; flip angle, 12°; matrix size, 256×256; FOV, 256mm; 176 slices; slice thickness, 1mm; voxel size, 1×1×1mm^3^). Structural proton density scans (TR, 2.7s; TE, 36ms; flip angle, 90°; matrix size, 512×512; FOV, 192mm; 39 slices; slice thickness, 2mm; voxel size, 0.37×0.37mm^3^) were also acquired prior to and in the same plane as each functional session in order to aid the co-registration of functional data with structural volumes, and identify the lateral geniculate nucleus. Functional data were acquired with a BOLD T2* EPI sequence (TR, 3s; TE, 30ms; flip angle 90°; matrix size, 128×128; FOV, 192mm; 39 slices; slice thickness, 2mm; voxel size, 1.5×1.5×2mm^3^).

Each participant underwent one high resolution structural scan, and two sessions consisting of a structural proton density scan followed by functional runs mapping the visual field (one expanding ring, one rotating wedge), testing each hemifield separately.

### Retinotopic Mapping

Visual maps that included the far peripheral representation were extracted using phase-encoded retinotopic mapping (DeYoe et al., 1996, Sereno et al., 1995, Engel et al. 1997). Visual stimuli were generated either with MatLab (version R2012a; The MathWorks, Natick, MA) or Psykinematix 1.4 (Beaudot, 2009) on a Mac Mini OS X and projected to participants through the scanner bore with either a Dukane 8942 ImagePro or PROPixx DLP LED projector. These included wedge and ring stimuli and both they contained high contrast reversing checker boards, flickering at a rate of 6 Hz (100% luminance contrast, 50% luminance background). The rotating wedge stimulus in both extended horizontally to 72°, vertically ca. 20° from fixation, covered 30° of the visual field. The MatLab generated stimulus stepped in 11.25° increments, flickering at 6 Hz, whilst the Psykinematix wedge stimulus stepped in 15° increments and flickered at a rate of 4 Hz, beginning at the upper vertical meridian within both hemifield conditions. The expanding annulus in both stimulus sets had a total annular width of 30°. As the rings approached the edge of the stimulated field, each ring was replaced by a new one originating from the centre of fixation. The Psykinematix generated rotating wedge stimulus beginning at the upper vertical meridian for the right hemifield, and at the lower vertical meridian in the left hemifield. A fixation cross, a grey ‘+’ sign, 0.87° in size, or a red ‘+’ sign, 0.6° in size was present throughout the entire scan. Participants viewed the stimuli at a distance of 275 mm from a supine position through a wide mirror mounted on the head coil, allowing them to see the projection on a custom in-bore acrylic screen (3050mm x 2030mm) mounted behind the head coil (see Figure 2.3). Mean luminance of the stimulus was 98 cd/m^2^. The head of each subject was stabilised with foam pads placed inside the head coil and a forehead velcro-strap to reduce motion artefacts. In order to increase the restricted field of view normally imposed by the MRI scanner, each hemifield was tested in separate functional scans allowing the stimulus to extend to 72° along the horizontal meridian. Participants viewed stimuli passively, continuously fixating a small cross placed either on the left or right side of the screen depending on which hemifield was being tested. Except in the first few sessions (4 hearing, 5 deaf), the head was tilted slightly (ca. 3°) towards the fixation cross to maximise participant comfort, field of view and fixation and head stability.. The cortical representation of polar angle as well as eccentricity (distance from centre of gaze) of the visual field were mapped for each participant. Deaf participants were shown instructions on the screen during the scanning session, and indicated their understanding by pressing a button. Hearing participants were spoken to via the built-in scanner intercom system.

### V1 Region-Of-Interest Measurements

The delineation of primary visual cortex, V1, was based on previous literature, which established the identifying features of visual areas (Engel et al., 1997; Sereno et al., 1995; DeYoe et al., 1996; Wandell et al., 2007). The sub-ROI borders were guided by the eccentricity (expanding ring) data. These ROIs were defined by eccentricity bins which would provide an approximately equal number of voxels in each, in order to make these comparable between groups. The ROIs will therefore not be equal in terms of visual field angle due to cortical magnification, and as each fMRI volume is acquired every 3 seconds (TR), the eccentricity stimulus is designed to expand in discrete steps of 3 seconds, to which the above V1 ROIs were matched to as well. These bins corresponded to the representations of central (0-15°), mid-peripheral (15-39°) and far-peripheral visual field (39-72°). Data representing the given response phase corresponding to the given extent of the visual field stimulated was displayed on the flattened cortex. This guided the manual selection of cortical surface devoted to representing the given representation, and the definition included all voxels, and was not restricted to the inclusion of only active voxels. The cortical volume (mm^3^), surface area (mm^2^) and grey matter thickness (mm) were extracted for each sub-ROI, within each visual area. The surface area measurements were made on the 3D cortical manifold, following the method used in Dougherty et al. (2003). In this method, the visual areas are outlined on a 2D flat map, then transformed into the 3D manifold. The surface area was calculated by taking the coordinates belonging to the selected ROI and finding the nearest node on the 3D manifold describing the boundary of grey and white matter.

### Cortical Magnification Estimates

The areal cortical magnification was estimated in V1 by dividing the cortical surface area for each ROI (central, mid-peripheral and far-peripheral) by the area of visual field represented (mm^2^/degrees^2^). As the areal cortical magnification function typically follows an inverse exponential (Daniel & Whitteridge, 1961), values were log transformed (log(mm^2^/degrees^2^) and fit with a linear regression. For each participant, linear fit gradients were calculated for left and right V1, then averaged across hemispheres and then averaged across participants for each group. This provided a metric which is not influenced by individual differences in visual area size (Andrews et al., 1997; Dougherty et al., 2003).

### Lateral Geniculate Nucleus

The left and right lateral geniculate nuclei were identified in each participant using high resolution proton density scans taken in the same slice orientation and location as functional scans (see MRI protocol section) for optimum visualisation of subcortical structures (Fujita et al, 2001; Dai et al., 2011 ; Gupta et al., 2008). Regions of interest were identified on anonymised scans (KY) and checked by a second individual (HB) such that the identifier was unaware of whether scans belonged to deaf or hearing participants. Functional voxels falling within the region of interest were analysed from expanding ring scans as with cortical voxels (FFT analysis, phase yields preferred retinal eccentricity). A coherence threshold of > 0.23 (uncorrected p<0.01) was applied to exclude noisy voxels. As the LGN is small, each participant only contributed a few voxels; therefore, voxels were accumulated across all participants within each group. Group histograms were plotted in 3 deg eccentricity bins based on the *histcount* function in Matlab, which picks optimal bin size based on the number and distribution of values. As distributions were skewed towards lower eccentricities, data were log transformed to normalise distributions before statistical evaluation.

### Data and statistical analysis

All the measures described above were tested statistically with R (R Core Team, 2019 and SPSS (IBM SPSS Statistics 20). All t-tests described in the work were Welch two sample t-test. Before conducting the general comparison of total V1 volume, a normal distribution confirmed by Shapiro-Wilk test, W= 0.9792, p = 0.776). Dai, H., Mu, K. T., Qi, J. P., Wang, C. Y., Zhu, W. Z., Xia, L. M., … Morelli, J. N. (2011).

